# Assembloid model to study loop circuits of the human nervous system

**DOI:** 10.1101/2024.10.13.617729

**Authors:** Yuki Miura, Ji-il Kim, Ovidiu Jurjuț, Kevin W. Kelley, Xiao Yang, Xiaoyu Chen, Mayuri Vijay Thete, Omer Revah, Bianxiao Cui, Marius Pachitariu, Sergiu P. Pașca

## Abstract

Neural circuits connecting the cerebral cortex, the basal ganglia and the thalamus are fundamental networks for sensorimotor processing and their dysfunction has been consistently implicated in neuropsychiatric disorders^1-9^. These recursive, loop circuits have been investigated in animal models and by clinical neuroimaging, however, direct functional access to developing human neurons forming these networks has been limited. Here, we use human pluripotent stem cells to reconstruct an *in vitro* cortico-striatal-thalamic-cortical circuit by creating a four-part loop assembloid. More specifically, we generate regionalized neural organoids that resemble the key elements of the cortico-striatal-thalamic-cortical circuit, and functionally integrate them into loop assembloids using custom 3D-printed biocompatible wells. Volumetric and mesoscale calcium imaging, as well as extracellular recordings from individual parts of these assembloids reveal the emergence of synchronized patterns of neuronal activity. In addition, a multi–step rabies retrograde tracing approach demonstrate the formation of neuronal connectivity across the network in loop assembloids. Lastly, we apply this system to study heterozygous loss of *ASH1L* gene associated with autism spectrum disorder and Tourette syndrome and discover aberrant synchronized activity in disease model assembloids. Taken together, this human multi-cellular platform will facilitate functional investigations of the cortico-striatal-thalamic-cortical circuit in the context of early human development and in disease conditions.

## Introduction

In the mammalian central nervous system, the corticostriatal-thalamic-cortical loop circuits connecting the cerebral cortex, basal ganglia, and thalamus are crucial for regulating movement, reward processing, and complex cognitive functions^1-10^. Disruptions in these circuits have been strongly associated with neurodevelopmental and neuropsychiatric conditions, including Tourette syndrome^11-13^. Although loop circuits have been extensively studied using clinical neuroimaging in human^14-17^ and animal models^1-10^, substantial gaps remain in our understanding of how these networks are established during human development and how activity patterns differ in the context of disease. These gaps are primarily attributed to the challenges in directly accessing patient brain tissue or difficulty in simultaneously probing activity across multiple brain regions in prenatal development.

Recent advances in stem cell-based neural assembloid technologies enabled direct aspects of human neural circuit development and have identified disease-associated phenotypes^18-26^. While two-part and three-part assembloids models have been successfully developed and more recently, a four-part assembloid in a linear configuration has been constructed^27^, the creation of loop assembloid remains unachieved.

In this study, we developed a human cellular model of the cortico-striatal-thalamic-cortical circuit *in vitro* by generating and assembling individual regionalized neural organoids that resemble the cerebral cortex that includes glutamatergic neurons, striatum that includes medium spiny neurons, mesencephalon that includes GABAergic projection neurons, and diencephalon that includes thalamic glutamatergic neurons. We designed a custom 3D well to functionally integrate these four neural organoids into loop-shaped assembloids (human loop assembloid, hLA). We implemented a custom volumetric and mesoscale calcium imaging, as well as extracellular recordings from individual components of these assembloids, and observed the gradual emergence of synchronized neuronal activity patterns in the hLA. Furthermore, multi-step rabies virus tracing revealed the formation of neuronal connectivity in the assembloids. Lastly, we demonstrate that this platform can be used to study cellular and network-level phenotypes of a human neurodevelopmental disorder. Specifically, following a heterozygous loss of the histone methyltransferase *ASH1L*, a gene identified as a high-risk factor for autism spectrum disorder (ASD) and Tourette syndrome, we identified altered neural synchronization in hLA.

## Results

### Generation of individual components of the cortico-striatal-thalamic-cortical circuit

To generate the key cell types of the cortico-striatal-thalamic-cortical circuit *in vitro*, we differentiated human induced pluripotent stem (hiPS) cells into regionalized neural organoids using specific small molecules and growth factors (**Fig. 1a, b, Supplementary Data Fig 1**). We followed our previously validated protocol ^20,28^ to generate human cortical organoids (hCO) and human diencephalic organoids (hDiO), which contain cortical glutamatergic neurons and thalamic glutamatergic neurons, respectively. To derive neural organoids that give rise to cells in parts of the basal ganglia, including striatal projection neurons (SPNs), we modified our previous protocol for generating human striatal organoids (hStrO) ^21,29^. Specifically, this modified protocol generates and includes expanded cell diversity in hStrO by modulating WNT, Sonic Hedgehog (SHH) pathways, and activating retinoic acid receptors which are highly expressed in the developing striatum^30^. Furthermore, we established a protocol to derive human mesencephalic organoids (hMeO) that contain GABAergic and dopaminergic neurons, modified from a bi-phasic WNT pathway activation protocol ^31^. To verify cell identity in these regionalized organoids, we performed droplet-based singlecell RNA sequencing (scRNA-seq) of individual hCO, hStrO, hMeO and hDiO (day 100-128) across 2 hiPS cell lines, and integrated a total of 32,408 high-quality cells (**Fig. 1c, d, Supplementary Data Fig. 2**). Regional mapping to a human fetal brain atlas ^32^ demonstrated that cells derived from each regionalized organoid resembled their corresponding *in vivo* counterparts. (**Fig. 1e, f**).

**Fig 1.**
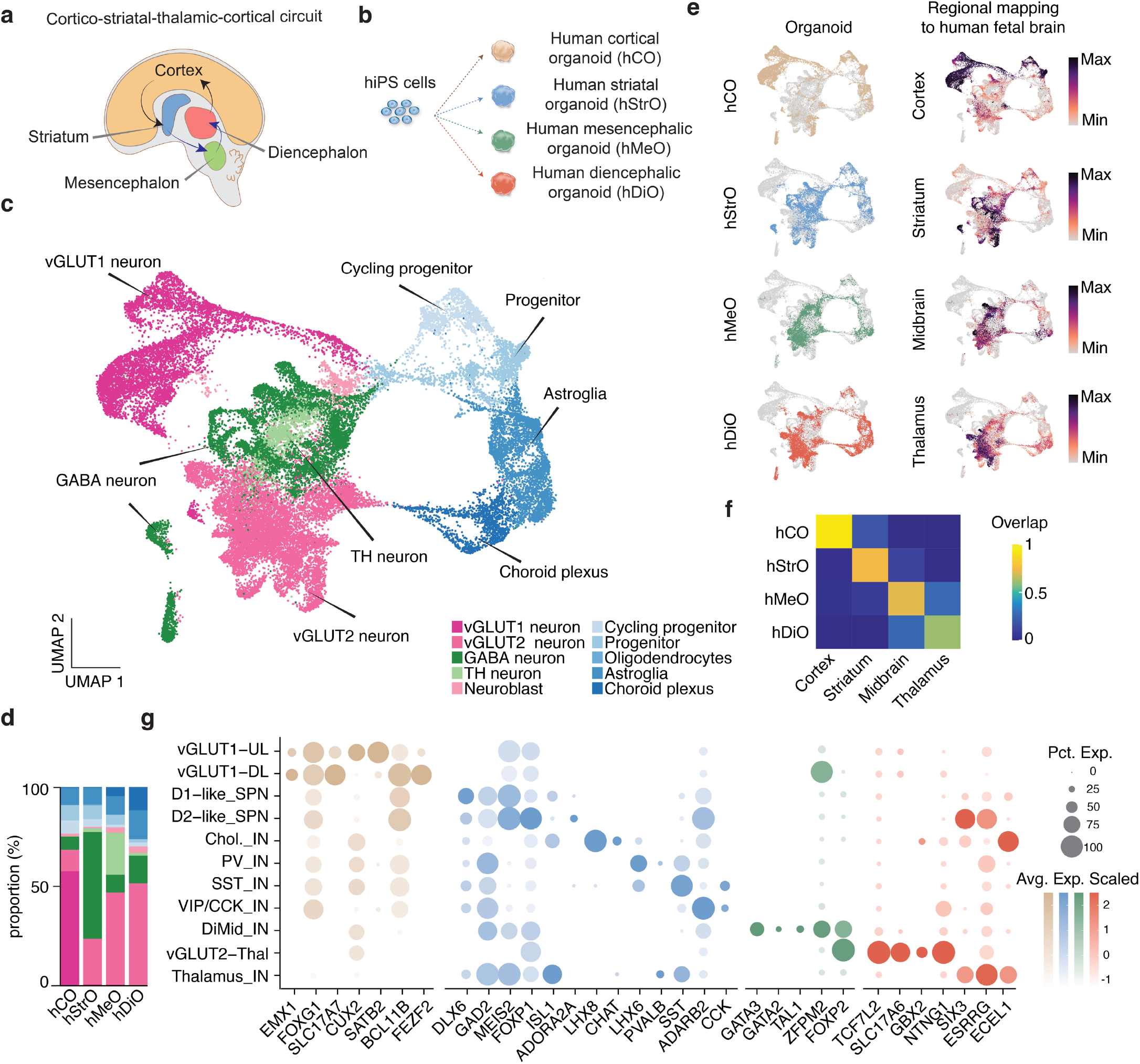
Generation of key cell types of the human cortico-striatal-thalamic-cortical circuit. **a**.Schematic illustrating the key components of the cortico-striatal-thalamic-cortical circuit. **b**. Schematic illustrating the generation of human cortical organoids (hCO), human striatal organoid (hStrO), human mesencephalic organoid (hMeO), and human diencephalic organoids (hDiO) from hiPS cells. **c**. UMAP visualization of single cell gene expression of hCO, hStrO, hMeO, and hDiO (n = 11452 cells for hCO day 100–128, n = 7986 cells for hStrO day 100, n = 5761 cells for hMeO day 100, n = 9009 cells for hDiO day 100 from 2 hiPS cell lines). **d**. Distribution of cell clusters across regional organoids, with color labels consistent with those shown in panel c. **e**. UMAP plots colored by the organoid identities (left) and regional mapping of the human fetal brain atlas for each region ^32^. **f**. Quantification of proportion of indicated human fetal brain region mapping to indicated regionalized organoid by transfer labeling to the reference human fetal brain atlas. **g**. Dot plot of selected marker gene expression in each single neuronal subcluster.

**Fig 2.**
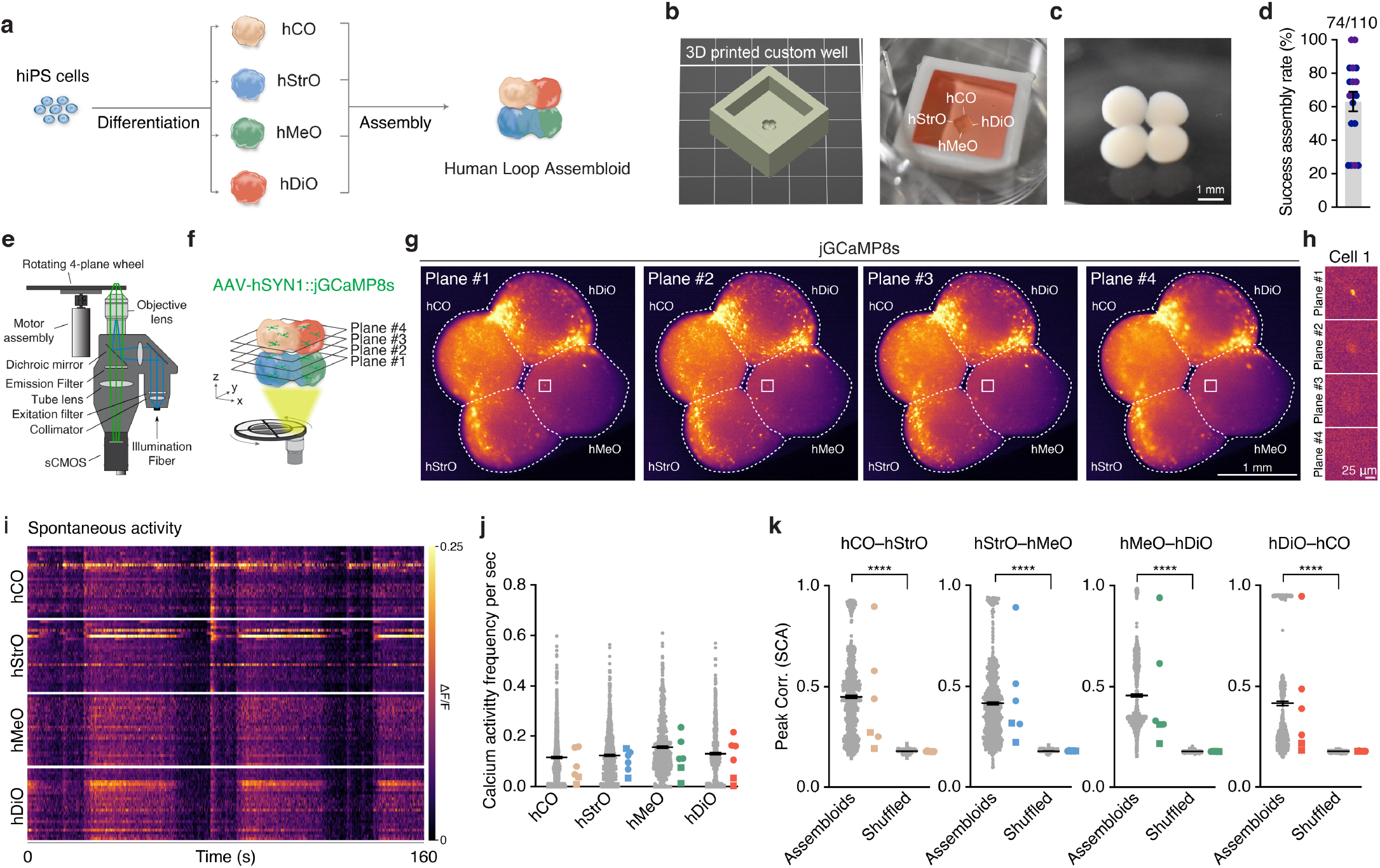
Generation and calcium imaging of human loop assembloids. **a**. Schematic illustrating the generation and fusion of hCO, hStrO, hMeO and hDiO to form a human loop assembloid. **b**. Schematic (left) and representative image (right) of a 3D printed biocompatible custom well designed to hold organoids in respective positions during assembly of loop assembloids. **c**. Representative image of a successfully assembled loop assembloid. **d**. Success assembly rate of loop assembloids from 18 batches, 9 differentiations, 2 hiPS cell lines. **e**. Schematic illustrating a widefield and multi-Z-plane imaging system with a rotating wheel on top of an objective lens. **f**. Schematic illustrating live calcium imaging of a loop assembloid captured from four focal planes, with 100 μm distance between planes, determined by different wheel positions.=\\ **g**. Representative images showing jGCaMP8s signals detected from four different Z planes within the loop assembloid. **h**. Representative images showing an example of a single cell that was captured in Plane #1 but not detected in other planes. **i**. Representative traces of calcium activity from 100 cells (25 from each region) detected throughout the entire loop assembloid. **j**. Quantification of the number of spontaneous calcium transients captured in different regions of the loop assembloid (n = 874 cells in hCO, n = 836 cells in hStrO, n = 740 cells in hMeO, n = 681 cells in hDiO from 6 assembloids, 2 hiPS cell lines in 3 differentiations) at day 146–162. **k**. Correlation values averaged per cell, for connections within loop assembloids, comparing to shuffled data (n = 874 cells in hCO–hStrO, n = 836 cells in hStrO–hMeO, n = 740 cells in hMeO–hDiO, n = 681 cells in hDiO–hCO from 6 assembloids, 2 hiPS cell lines in 3 differentiations, Wilcoxon test, \*\*\*\**P* < 0.0001) at day 146–162. Data is shown as mean ± s.e.m. Dots indicate batches in d. Small dots indicate units or cells and large dots indicate assembloids in j, k. Each shape represents a hiPS cell line: Circle, 1205-4; Rectangle, KOLF2.1J in j, k.

Analysis of the integrated dataset revealed distinct clusters corresponding to cortical glutamatergic neurons (VGLUT1/*SLC17A7*^+^, *FOXG1*^+^, *EMX1*^+^), striatal GABAergic neurons (*GAD1*^+^, *GAD2*^+^, *FOXG1*^+^, *DLX6*^+^), mesencephalon dopaminergic neurons (*EN1*^+^, *TH*^+^, *SLC6A3*^+^), and thalamic glutamatergic neurons (VGLUT2/*SLC17A6*^+^, *TCF7L2*^+^) (**Fig. 1c, Table S1**). Subclustering of GABAergic neurons revealed a diversity of SPN populations (D1-like SPN: *MEIS2*^+^, *FOXP1*^+^, *ISL1*^+^; D2-like SPN: *MEIS2*^+^, *FOXP1*^+^, *ADORA2A*^+^), in addition to putative striatal interneurons (cholinergic interneurons: *CHAT*^+^, *LHX8*^+^; PV interneurons: *PVALB*^+^, *LHX6*^+^; somatostatin INs: *SST*^+^, *LHX6*^+^; CCK/VIP INs: *CCK*^+^, *ADARB2*^+^) (**Fig. 1g, Supplementary Data Fig. 2, Table S2**). Furthermore, we identified clusters of GABAergic neurons resembling thalamic reticular nucleus (*SIX3*^+^, *ISL1*^+^, *PVALB*^+^) ^33^ derived from both hDiO and hStrO samples as well as GABAergic neurons in hDiO and hMeO that mapped to the mesencephalon and expressed markers of the developing substantia nigra pars reticulata (*GATA3*^+^, *GATA2*^+^, *TAL1*^+^, *ZPFM2*^+^)^34^. Taken together, the transcriptomic analysis demonstrates that regionalized organoids encompass critical cellular components of the human loop circuits, including glutamatergic neurons of the cortex and thalamus as well as GABAergic neurons of the striatum and mesencephalon.

### Constructing loop assembloids and probing calcium activity with widefield, multi-Z-plane imaging

We next aimed to assemble these regionalized organoids as arranged in a loop configuration that comprise the four organoids described above: hCO, hStrO, hMeO, and hDiO in the right order and positions (**Fig. 2a**). Unlike the linear four-part assembloid we recently reported^27^, this circular orientation was prone to disassembly and challenging to maintain. Therefore, we designed a biocompatible, custom 3D-printed well (**Fig. 2b**). Using this inexpensive, custom well as an assembly scaffold, we achieved precise placement of the four regionalized organoids. After 4–7 days, organoids integrated into a stable, loop-shaped assembloid that we refer to as human loop assembloids (hLA) (**Fig 2c**). We evaluated the success rate of achieving the desired orientation and stable assembly and found that approximately 60% of the samples were successfully fused and maintained their assembly when using the 3D-printed well (**Fig. 2d**).

After assembly, hLA can span up to 4 mm in maximum diameter. To monitor neuronal activity in this entire large multi-cellular 3D structure, we built a custom imaging system with a broad field-of-view that also captures calcium activity at single-cell resolution across multiple Z-planes (**Fig. 2e and f, Supplementary Data Fig 3a and b**). To achieve accurate and high-speed imaging (0.5–20 Hz) across four Z-planes, we incorporated a rotating wheel with four different thicknesses for light reflection (**Fig. 2e, f**). This wheel rotated during image acquisition and changed the focal plane in 100 μm increments allowing imaging of different focal planes of the assembloid (**Supplementary Data Fig. 3b**). We validated uniform detection with minimal simulated distortion values across the wide field-of-view (**Supplementary Data Fig. 3c**), and confirmed the consistency of this high-speed, four Z-plane imaging (5 Hz) across over a thousand frames (**Supplementary Data Fig. 3d**).

**Fig 3.**
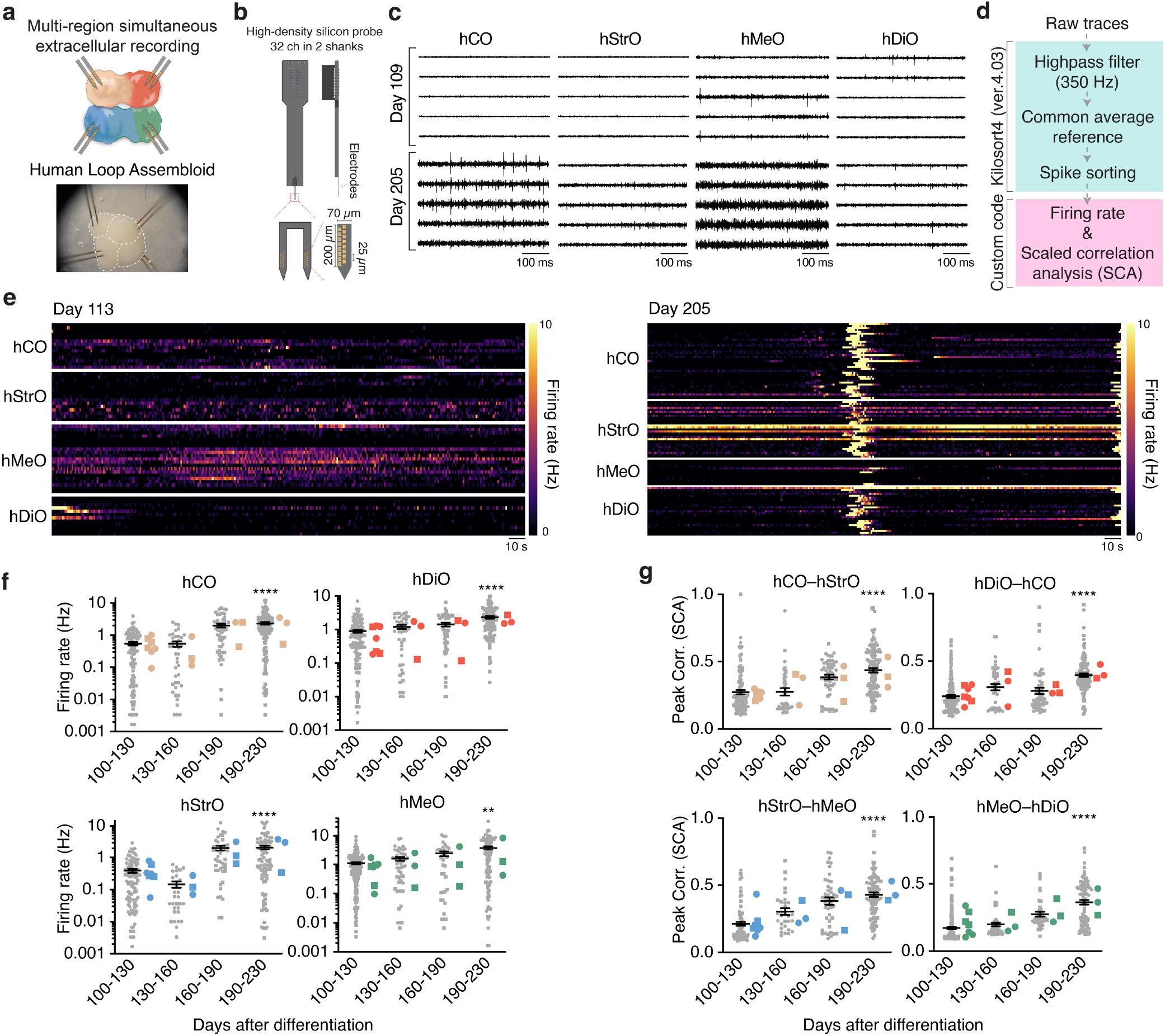
Multi-regional extracellular recordings of human loop assembloids. **a**. Schematic illustrating the setup for multi-regional simultaneous extracellular recordings in a loop assembloid (top) and a representative image showing the placement of silicon probes within a loop assembloid during recordings (bottom). **b**. Schematic illustrating the design of a high-density silicon probe with 32 channels distributed across 2 shanks. **c**. Representative raw traces of neuronal activity recorded from each region of a loop assembloid (hCO, hStrO, hMeO, and hDiO) at day 109 and day 205. **d**. Schematic of the data processing pipeline used to analyze the recorded signals. **e**. Heatmaps showing the neuronal activity recorded from each region of loop assembloids at day 113 (left) and day 205 (right). Each row represents one sorted unit. **f**. Quantification of firing rates over time in each region of loop assembloids (hCO; *n* = 98 units for days 100–130, *n* = 41 units for days 130–160, *n* = 54 units for days 160–190, *n* = 124 units for days 190–230 / hStrO; *n* = 91 units for days 100–130, *n* = 30 units for days 130–160, *n* = 46 units for days 160–190, *n* = 88 units for days 190–230 / hMeO; *n* = 183 units for days 100–130, *n* = 51 units for days 130–160, *n* = 38 units for days 160– 190, *n* = 86 units for days 190–230 / hDiO; *n* = 134 units for days 100–130, *n* = 43 units for days 130–160, *n* = 50 units for days 160–190, *n* = 113 units for days 190–230, from 7 assembloids for days 100–130, 3 assembloids for days 130–160, 3 assembloids for days 160–190, 3 assembloids for days 190–230, 2 hiPS cell lines in 5 differentiations, Kruskal-Wallis test with Dunn’s multiple comparisons test, \*\**P* = 0.0011, \*\*\*\**P* < 0.0001). **g**. Correlation values averaged per cell, for connections within loop assembloids (hCO–hStrO; *n* = 97 units for days 100–130, *n* = 36 units for days 130–160, n = 54 units for days 160–190, n = 123 units for day 190–230 / hStrO–hMeO; n = 90 units for day 100–130, *n* = 30 units for day 130–160, n = 44 units for day 160–190, n = 88 units for day 190–230 / hMeO–hDiO; n = 183 units for day 100–130, n = 49 units for day 130–160, n = 38 units for day 160–190, n = 85 units for day 190–230 / hDiO–hCO; n = 134 units for day 100–130, n = 42 units for day 130–160, n = 50 units for day 160– 190, n = 112 units for day 190–230, from 7 assembloids for day 100–130, 3 assembloids for day 130–160, 3 assembloids for day 160–190, 3 assembloids for day 190–230, 2 hiPS cell lines in 5 differentiations, Kruskal-Wallis test with Dunn’s multiple comparisons test, \*\*\*\**P* < 0.0001). Data is shown as mean ± s.e.m. Small dots indicate units and large dots indicate assembloids in f, g. Each shape represents a hiPS cell line: Circle, 1205-4; Rectangle, KOLF2.1J in f, g.

To probe neuronal calcium activity in hLAs, we virally expressed jGCaMP8s under the hSYN1 promoter^35^. This enabled detection of calcium activity across all four regions of the hLA in four Z-planes at single-cell resolution at day 146 (**Fig. 2g**), capturing cells that would have been missed with single-plane imaging (**Fig 2h**). We extracted spontaneous calcium activity from hundreds to thousands of cells simultaneously in hLAs (**Fig 2i**). (**Fig 2j**). Simultaneous probing of calcium activity in all four hLA regions allowed the exploration of collective patterns of activity that may emerge from their interactions. Remarkably, we found that neurons within hLAs demonstrated synchronous activity across all four regions at day 146–162 (**Fig 2i**). We quantified correlation indices using scaled correlation analysis and revealed significantly higher correlation values in hLA compared to shuffled datasets across all four regions (**Fig. 2k**), indicating coordinated activity across the four parts of the hLA.

We further performed multi-region extracellular recordings of spontaneous activity in the hLA using multiple high-density silicon probes with 128 total channels over 4 probes with 2 shanks each (**Fig. 3a, b**). Inspection of raw traces at multiple time points of hLA revealed spikes in multiple channels across the four regions (**Fig 3c**). To analyze the activity of single units, we applied Kilosort 4 for spike sorting^36^. We extracted firing rates for each unit and calculated scaled correlations (**Fig 3d**). Interestingly, longitudinal measurements of activity showed that firing rates rise over time and became ∼2–5 fold higher in the late stage (day 190–230) compared to the early stage (day 100–130) of hLA (**Fig 3e, f**). Moreover, correlation values from inter-regional connectivity demonstrated a gradual increase over time (**Fig 3g**), indicating more synchronized activity patterns at the later stage of assembloids. In summary, we successfully generated hLAs using 3D-printed custom scaffolds and identified an emergent synchronized neuronal activity pattern over time in the assembloids, demonstrating that the assembly of regionalized organoids resembling the human cortex, striatum, midbrain, and thalamus enables the progressive formation of networks capable of synchronized neuronal activity.

### Retrograde viral tracing of projections in the hLAs

To verify projections within the loop circuit in the hLA, we employed rabies viral retrograde tracing to label projecting cells, as previously applied in the assembloid systems ^18,20,21,27^. First, we verified the projection from hCO to hStrO by infecting hStrO with a G-deleted rabies virus expressing Cre recombinase and the G protein under the EF1α promoter. Simultaneously, we infected hCO with AAV-DJ-EF1α::DIO-tdTomato before assembly ^37,38^. Retrograde transmission of the rabies virus would result in tdTomato expression in hCO neurons following Cre recombinase, and this would indicate some level of hCO to hStrO connectivity. Consistent with our previous report ^21^, we found tdTomato^+^ cells in the hCO side of the hCO–hStrO assembloid, indicating successful projections from the hCO to the hStrO. We also applied this strategy to other pairs in the hLA and confirmed retrograde labeling from hStrO to hMeO, from hMeO to hDiO, and from hDiO to hCO (**Fig. 4a**).

**Fig 4.**
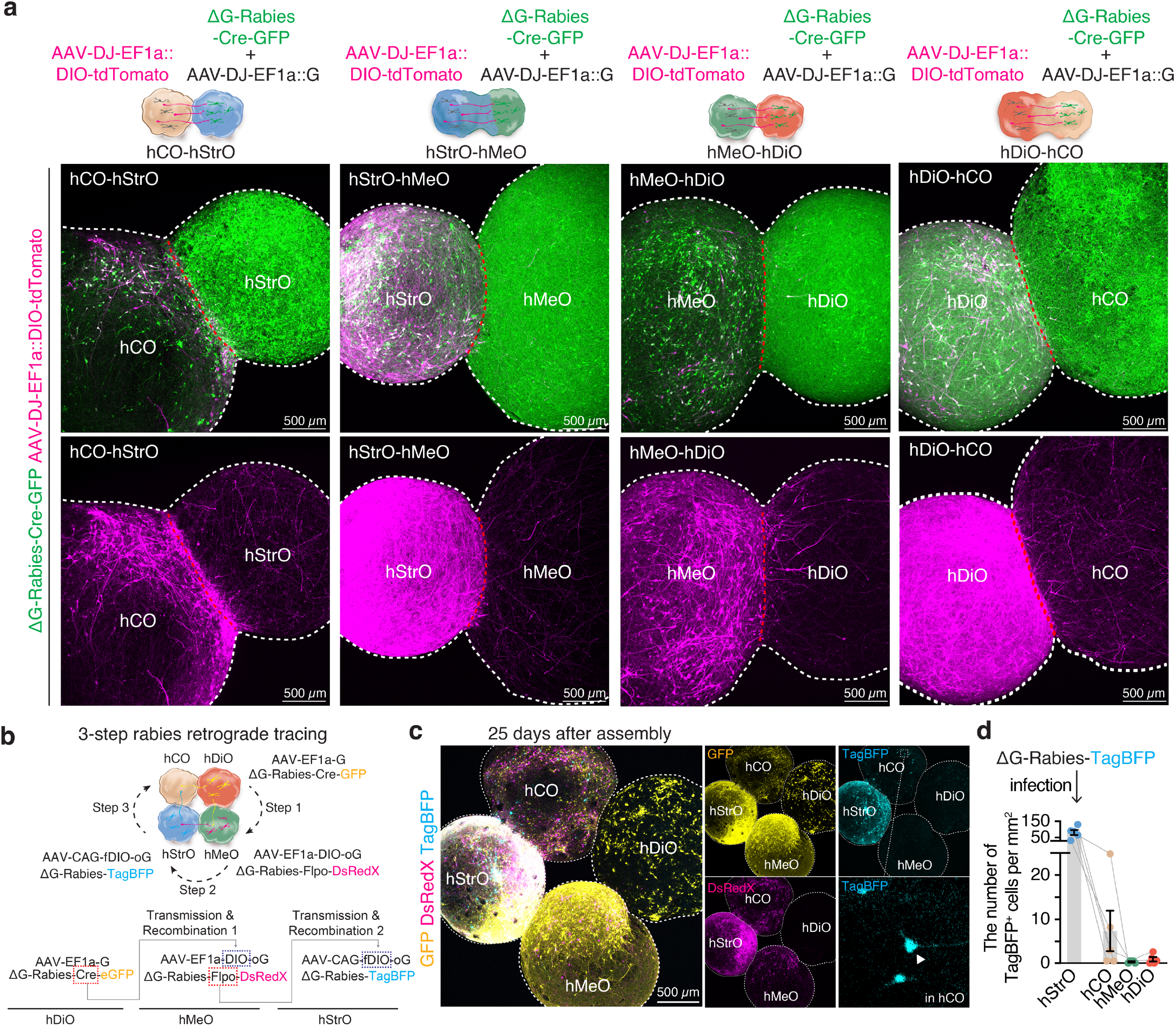
Retrograde projection tracing in human loop assembloids. **a**. Schematics (top) illustrating retrograde tracing in 2–part assembloids (hCO–hStrO, hStrO–hMeO, hMeO–hDiO and hDiO–hCO) using rabies virus and representative images (bottom) showing GFP and tdTomato expression. High-intensity images display strong tdTomato expression and their projections in the target regions. **b**. Schematics illustrating a 3–step rabies retrograde tracing strategy to trace loop projections from hDiO to hMeO to hStrO to hCO. This strategy utilized Cre and Flpo recombinases and DIO– and fDIO– forms of oG, necessary for the transmission of rabies virus, to restrict transmission to neurons where Cre or Flpo recombinases are present. **c**. Representative images showing the results of 3–step rabies retrograde tracing in a loop assembloid. Insets highlight TagBFP expression in the hCO part of the loop assembloid, indicating loop connectivity and successful retrograde transmission of the rabies virus. **d**. Quantification of the number of TagBFP+ cells per mm^2^ in each region of loop assembloids (n = 6). Data is shown as mean ± s.e.m. Dots indicate assembloids in d.

Next, we implemented a three–step and multi–color rabies virus retrograde tracing strategy to examine loop connectivity within hLAs (**Fig. 4b**). The combination of rabies viruses expressing different recombinases (Cre or Flpo) and double floxed inverted (DIO– or fDIO–) optimized rabies glycoprotein (oG) should enable tracing of up to 3 connections across 4 components (**Fig 4b**). Expression of the rabies glycoprotein (G) under the EF1α-promoter would enable the transmission of rabies virus carrying Cre–EGFP from hDiO to hMeO. In hMeO, DIO-oG would be inverted and expressed by Cre recombinase, allowing transmission of Flpo–DsRedX to hStrO. Subsequently, Flpo recombinase would invert fDIO-oG and enable transmission into hCO of rabies virus containing Tag-BFP in hStrO. Therefore, the rabies viral transmission would take place across the loop, and this would result in TagBFP expression in hCO in the assembloids. Indeed, in recombination experiments performed in 6 hLA, we detected TagBFP^+^ cells in the hCO part of assembloids (**Fig. 4c, d**), indicating successful 3-step retrograde transmission that may underlie the synchronized neural activity we observed by imaging and extracellular recordings.

### Altered neural activity in *ASH1L* KO assembloids

Lastly, we verified the ability of hLAs to identify cellular or circuit level phenotypes associated with a disease risk gene. Gene variants in the histone methyltransferase *ASH1L* have been linked to ASD^39,40^, epilepsy and Tourette syndrome^41^. We first confirmed that *ASH1L* is widely expressed across the majority of cell types in the four types of neural organoids (**Supplementary Data Fig. 4a**). To model haploinsufficiency of *ASH1L*, we used CRISPR–mediated gene editing to generate heterozygous *ASH1L* knockout (*ASH1L*^+/-^) hiPS cell lines (**Fig. 5a**, **Supplementary Data Fig. 4b**), and confirmed the heterozygous, one nucleotide insertion within the gRNA target region by Sanger sequencing (**Fig. 5 b, c**). We then generated four regionalized organoids from isogenic control and *ASH1L*^+/-^ and found that the size of *ASH1L*^+/-^ hStrO was smaller compared to control (**Supplementary Data Fig. 4c, d**), while other neural organoids were unaffected. We assembled these organoids to generate hLAs from control and *ASH1L*^+/-^ hiPS cell lines to explore the circuit–level consequences of *ASH1L* haploinsufficiency. Simultaneous extracellular recordings from each region of assembloids revealed that *ASH1L*^+/-^ hLAs exhibited highly repeated and synchronized neural activity patterns across all regions in the assembloids (**Fig. 5d**, **Supplementary Data Fig. 5a, b**). Firing rates were significantly increased in hCO, hMeO, hDiO (**Fig 5e**), while scaled correlation coefficients were higher in all pairs of organoids in *ASH1L*^+/-^ hLAs (**Fig 5f**). This is consistent with a neuroimaging study from patients in Tourette’s syndrome, showing increased basal ganglia-cortical and thalamo-cortical connectivity ^11^. Using bubble plots, we visualized both spiking activity and connectivity across the loop, In the bubble plot, lines between organoids or around an individual organoid indicate inter– and intra– regional correlation respectively, while the size of the circle indicates firing rates (**Fig 5g**). Interestingly, only hStrO that were reduced in size in *ASH1L*^+/-^ showed unaltered firing rates, while all the other regions were hyperactive. These results highlight the capability of hLA to capture cellular and circuit level dysfunction associated with haploinsufficiency of *ASH1L* and thus to model aberrations in emergent properties of the human cortico-striatal-thalamiccortical circuit.

**Fig 5.**
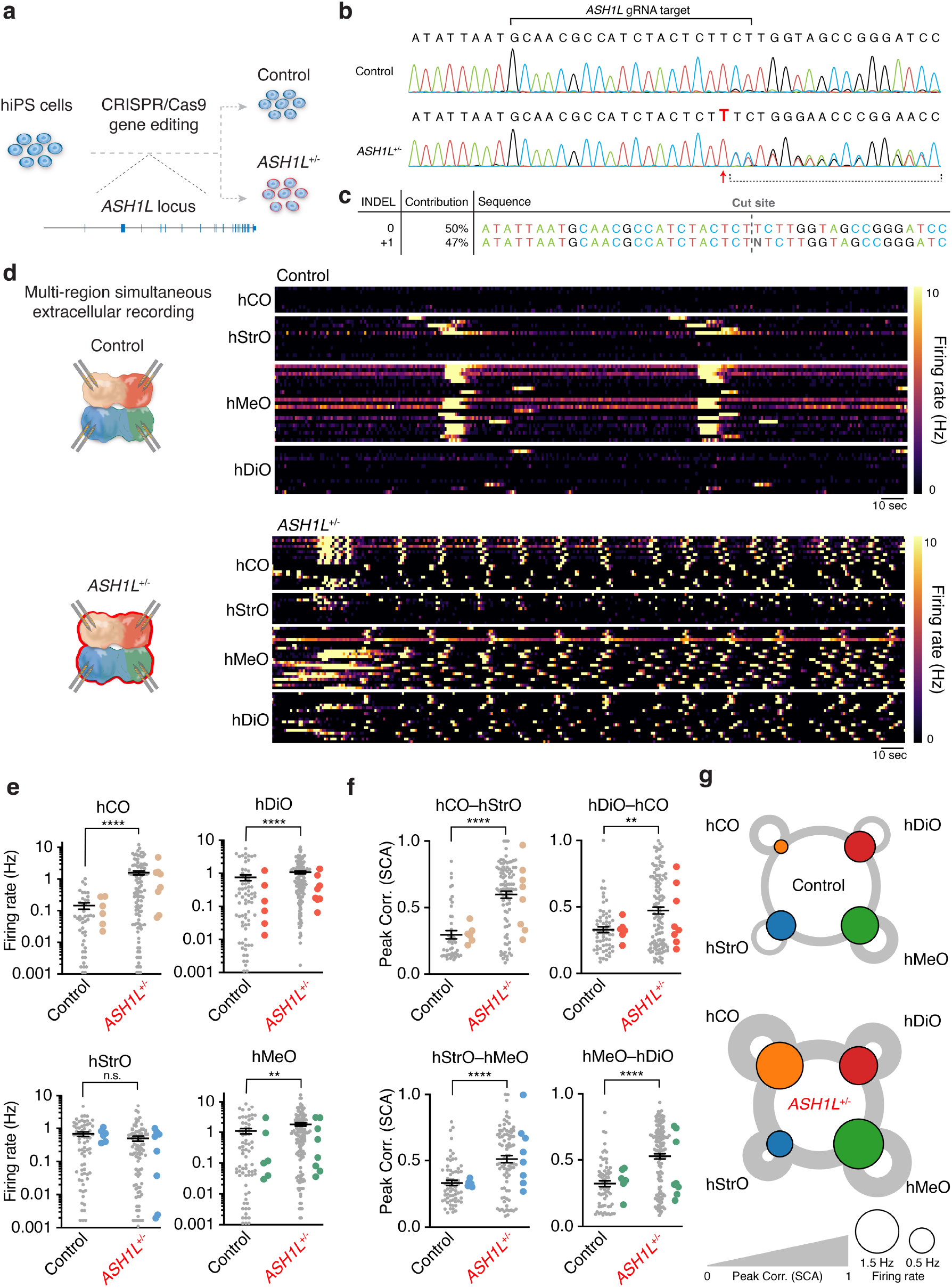
Human loop assembloid model of *ASH1L* haploinsufficiency. **a**. Schematic illustrating the strategy used to generate *ASH1L*^+/-^ hiPS cells using CRISPR/Cas9 gene editing. **b**. Sanger sequencing results of control hiPS cells (top) and genome edited *ASH1L*^+/-^ hiPS cells (bottom). **c**. Allelic frequency of INDELs based on ICE analysis on the Sanger sequencing results. **d**. Schematic illustrating multi-region simultaneous extracellular recordings (left) and heatmaps showing neuronal activity from control and *ASH1L*^+/-^ assembloids. Each row represents one sorted unit. **e**. Quantification of firing rates in each region of control and *ASH1L*^+/-^ loop assembloids (hCO; n = 45 units for control, n = 107 units for *ASH1L*^+/-^ / hStrO; n = 69 units for control, n = 88 units for *ASH1L*^+/-^ / hMeO; n = 69 units for control, n = 139 units for *ASH1L*^+/-^ / hDiO; n = 69 units for control, n = 125 units for *ASH1L*^+/-^, 6 control assembloids and 8 *ASH1L*^+/-^ assembloids from 4 differentiations, Mann-Whitney test, n.s. = not significant, \*\**P* = 0.0020, \*\*\*\**P* < 0.0001). **f**. Correlation values averaged per cell, for connections within control and *ASH1L*^+/-^ loop assembloids (hCO–hStrO; n = 44 units for control, n = 92 units for *ASH1L*^+/-^ / hStrO–hMeO; n = 66 units for control, n = 82 units for *ASH1L*^+/-^ / hMeO–hDiO; n = 65 units for control, n = 133 units for *ASH1L*^+/-^ / hDiO–hCO; n = 63 units for control, n = 118 units for *ASH1L*^+/-^, 6 control assembloids and 8 *ASH1L*^+/-^ assembloids from 4 differentiations, MannWhitney test, n.s. = not significant, \*\**P* = 0.0021, \*\*\*\**P* < 0.0001). **g**. Bubble plots representing the average firing rates in each region and the correlation values across all samples for control (top) and *ASH1L*^+/-^ (bottom) assembloids. Data is shown as mean ± s.e.m. Small dots indicate units and large dots indicate assembloids in e, f.

## Discussion

Modeling human brain loop circuits *in vitro* presents a powerful opportunity to explore mechanisms underlying the early development of the human cortico-striatal-thalamiccortical circuit and to decipher the pathophysiology of the disease. In previous work, we demonstrated that assembloids composed of regionalized neural organoids derived from hiPS cells could effectively model the formation of human neural circuits, including responses to noxious stimuli and muscle contractions induced by cortical neuron stimulation^18-21,27^. In this study, we extended this approach to construct a four-component loop neural circuit, beginning with cortical neurons and connecting through striatal, midbrain, and thalamic neurons before returning to the cortical region. This human loop assembloid system represents an initial effort to model closed–loop pathways. To investigate neural activity within this system, we implemented volumetric and mesoscale calcium imaging alongside extracellular recordings, enabling simultaneous assessment of neural activity across the entire loop assembloid. Using advanced analysis pipelines, we extracted firing rates and estimated intra– as well as inter– regional connectivity, revealing the temporal progression towards synchronized patterns of activity. The formation of neuronal connectivity was corroborated by a multi-step rabies retrograde tracing approach in the assembloids. Furthermore, we discovered that haploinsufficiency of *ASH1L* led to increased synchronous activity across the circuit, suggesting that abnormal activity patterns throughout the entire loop circuit may contribute to symptoms of *ASH1L*-associated neurodevelopmental disorders, such as Tourette syndrome. Animal models of *Ash1l* haploinsufficiency have been generated ^42,43^, but the functional simultaneous probing of the cortico-striatal-thalamic-cortical loop circuit has not been explored to date.

Improvements at several levels could further increase the utility of hLA. First, since hiPS cell-derived neurons are at prenatal or early postnatal stages even after hundreds of days of culture *in vitro*, improving maturation progression and scaling up the approach would expand the potential of this human multi-cellular model. Second, recent work has identified a more nuanced nature of classic loop circuits that involve other regions ^2^, including the direct connections from the thalamus to the striatum ^44^. We anticipate that at least some of these additional circuits could be constructed in parallel with hLA or within the same system. However, the implementation and usefulness of these *in vitro* models for disease modeling and drug screening will depend on keeping them as simple as possible, thus there are several advantages to applying these simplified neural circuit systems to unravel disease features. Third, further investigations are needed to explore the specificity of connectivity in these assembloids and the role of synchronized activity in establishing and refining these neural networks. These efforts could be directed towards uncovering the intrinsic principles underlying connectivity in early human circuits and developing strategies to accelerate the activity-dependent maturation of these *in vitro* models.

Overall, this new platform holds promise for providing insights into how human brain loop circuits are assembled and for advancing the screening of therapeutic compounds or the testing of genetic variants linked to neuropsychiatric disorders.

## Acknowledgements

We thank members of the Pasca laboratory at Stanford University for scientific inputs and the Stanford Wu Tsai Neurosciences Virus Core for producing AAVs. This work was supported by the Stanford Brain Organogenesis Big Idea Grant from the Wu Tsai Neurosciences Institute (to S.P.P.), the Wellcome leap 1kD program (to S.P.P.), National Institute of Mental Health (no. R01 MH115012) (to S.P.P.), Aligning Science Across Parkinson’s through the Michael J. Fox Foundation for Parkinson’s Research (MJFF) (to S.P.P.), the Kwan Funds (to S.P.P.), the Senkut Funds (to S.P.P.), the Coates Foundation (to S.P.P.), the Ludwig Family Foundation (to S.P.P.) the TAA Young Investigator Award (to Y.M.), the Stanford Maternal & Child Health Research Institute (MCHRI) Postdoctoral Fellowship (to Y.M. and O.R), and the National Research Foundation of Korea (to J.-i.K).

## Author contributions

Y.M., J.-i.K. and S.P.P. conceived the project and designed experiments. Y.M. developed the protocol to generate hStrO and hMeO, performed differentiation experiments, single-cell transcriptomics analyses, and functional imaging. J.-i.K. developed and carried out the multi-region extracellular recording. O.J. and M.P. developed custom analysis pipelines for calcium imaging and extracellular recordings. X.Y. and B.C. contributed to the design of the 3D custom wells. X.C. conducted CRISPR/Cas9 genome editing for *ASH1L* KO hiPS cell lines. M.V.T. contributed to the characterization of regionalized organoids. O.R. contributed to developing the custom microscope and extracellular recording platforms. Y.M., J.-i.K. and S.P.P. wrote the manuscript with input from all authors.

## Materials and Methods

### Characterization and maintenance of hiPS cells

In this study, human induced pluripotent stem (hiPS) cell lines were maintained and validated as standardized methods, as described previously^29,45,46^. Genome-wide SNP array was performed using Illumina genome-wide GSAMD-24v2-0 SNP microarray at the Children’s Hospital of Philadelphia (CHOP). Cell cultures were tested for and maintained Mycoplasma-free. Approval for this study was obtained from the Stanford IRB panel and informed consent was obtained from all subjects.

### Generation of hCO, hStrO, hMeO and hDiO from hiPS cells

To generate regionalized neural organoids, hiPS cells were cultured on vitronectin-coated plates (5 μg ml^-1^, Thermo Fisher Scientific, A14700) in Essential 8 medium (Thermo Fisher Scientific, A1517001). Cells were passaged every 4–5 days with UltraPure™ 0.5 mM EDTA, pH 8.0 (Thermo Fisher Scientific, 15575). For the generation of regionalized neural organoids, hiPS cells were incubated with Accutase^®^ (Innovative Cell Technologies, AT104) at 37°C for 7 min and dissociated into single cells. Optionally, 1–2 days before organoid formation, hiPS cells were exposed to 1% dimethylsulfoxide (DMSO) (Sigma-Aldrich, 472301) in Essential 8 medium. For aggregation into organoids, approximately 3 × 10^6^ hiPS cells were added per AggreWell-800 well in Essential 8 medium supplemented with the ROCK inhibitor Y27632 (10 μM, Selleckchem, S1049), centrifuged at 100 *g* for 3 min, and then incubated at 37°C in 5 % CO_2_. On the next day (day 1), organoids consisting of approximately 10,000 cells were collected from each microwell by pipetting medium in the well up and down with a cut P1000 pipet tip and transferred into ultra-low attachment plastic dishes (Corning, 3262) in Essential 6 medium (Thermo Fisher Scientific, A1516401) supplemented with patterning molecules as illustrated in **Supplementary Data Data Fig. 1**.

hCO were generated as previously described ^28,47,48^. For day 1 to 6, Essential 6 medium was changed every other day and supplemented with dorsomorphin (2.5 μM, Sigma-Aldrich, P5499) and SB-431542 (10 μM, R&D Systems, 1614) and was changed every day. On day 7, organoids were transferred to neural medium containing Neurobasal™-A Medium (Thermo Fisher Scientific, 10888022), B-27™ Supplement, minus vitamin A (Thermo Fisher Scientific, 12587010), GlutaMAX™ Supplement (1:100, Thermo Fisher Scientific, 35050079), Penicillin-Streptomycin (1:100, Thermo Fisher Scientific, 15070063), supplemented with FGF2 (20 ng/mL, R&D Systems, 233-FB) and EGF (20 ng/mL, R&D Systems, 236-EG) until day 22. From day 23 to day 46, the neural media was supplemented with BDNF (20 ng/mL, PeproTech, 450-02), NT3 (20 ng/mL, PeproTech, 450-03), L-Ascorbic Acid 2-phosphate Trisodium Salt (AA; 200 μM, Wako, 323-44822), N6, 2’-ODibutyryladenosine 3’,5’-cyclic monophosphate sodium salt (cAMP; 50 μM, Millipore Sigma, D0627), and cis-4, 7, 10, 13, 16, 19-Docosahexaenoic acid (DHA; 10 μM, Millipore Sigma, D2534). From day 47, neural medium containing B-27 Plus Supplement (Thermo Fisher Scientific, A3582801) was used for medium changes every 4 days.

hStrO were generated by a further optimized protocol based on our previous protocol^21^. For days 1 to 6, the Essential 6 medium was changed every other day and supplemented with dorsomorphin, SB-431542, and XAV-939 (1.25 μM, Tocris, 3748). On day 7, organoids were transferred to the neural medium. From day 7 to 17, the WNT pathway inhibitor IWP-2 (2.5 μM, Selleck Chemicals, S7085) was supplemented. From day 7 to 22, SAG (10 nM, Millipore Sigma, 566660-1MG) was supplemented. From day 13 to 23, Activin A (50 ng/mL, PeproTech, 120-14P) was supplemented, in addition to the compounds described above. From day 17 to 23, the retinoid X receptor agonist SR11237 (100 nM, Tocris, 3411) and the selective RARβ agonist CD2314 (100 nM, Tocris, 3824) were further supplemented. From day 23 to 46, the neural medium was supplemented with BDNF, NT3, AA, cAMP, and DHA. From day 43-46, neural spheroids cultures were supplemented with DAPT (2.5 μM, Stemcell Technologies, 72082) in addition to BDNF, NT-3, AA, cAMP and DHA. From day 47, medium containing B-27 Plus Supplement (Thermo Fisher Scientific, A3582801) and AA (200 μM) was used for medium changes every 4 days.

To generate hMeO, Essential 6 medium was changed every other day and supplemented with dorsomorphin and SB-431542 for days 1 to 6, and on day 7, organoids were transferred to the neural medium. From day 5 to 12, SAG (1 μM) was supplemented, and the WNT pathway activator, CHIR was supplemented as 3 μM from day 5 to 7, 7.5 μM from day 7 to 16, and 3 μM from day 17 to 22. From day 23 to 46, the neural medium was supplemented with BDNF, NT3, AA, cAMP, and DHA. For days 43–46, DAPT (2.5 μM) was additionally added. From day 47, medium containing B-27 Plus Supplement (Thermo Fisher Scientific, A3582801) and AA (200 μM) was used for medium changes every 4 days.

hDiO were generated as previously described^20^. For days 1 to 6, the Essential 6 medium was changed every day and supplemented with dorsomorphin and SB-431542. On day 5, the medium was additionally supplemented with 1 μM CHIR. On day 7, organoids were maintained in neural media supplemented with CHIR. On day 9 of differentiation, the neural medium was supplemented with 100 nM SAG (days 9–15), in addition to the CHIR. Furthermore, on days 12–18 of differentiation, organoids were supplemented with 30 ng/mL BMP7 (PeproTech, 120-03P), in addition to the compounds described above. From day 19 to 46, the neural medium was supplemented with BDNF, NT3, AA, cAMP, and DHA. For day 19–25, DAPT (2.5 μM) was additionally supplemented. From day 47, medium containing B-27 Plus Supplement (Thermo Fisher Scientific, A3582801) was used for medium changes every 4 days.

### Cryosection and immunocytochemistry

Organoids and assembloids were fixed in 4% paraformaldehyde (PFA)/phosphate buffered saline (PBS) overnight at 4°C. They were then washed in PBS and transferred to 30% sucrose/PBS for 2–3 days until the organoids/assembloids sank into the solution. Subsequently, they were rinsed in optimal cutting temperature (OCT) compound (Tissue-Tek OCT Compound 4583, Sakura Finetek) and 30% sucrose/PBS (1:1), embedded, and snap-frozen using dry ice. For immunofluorescence staining, 30 μm-thick sections were cut using a Leica Cryostat (Leica, CM1860). Cryosections were washed with PBS to remove excess OCT from the sections and blocked in 10% Normal Donkey Serum (NDS, Millipore Sigma, S30-100ML), 0.3% Triton X-100 (Millipore Sigma, T9284-100ML), and 1% BSA diluted in PBS for 1 hour at room temperature. The sections were then incubated overnight at 4°C with primary antibodies diluted in PBS containing 2% NDS and 0.1% Triton X-100. PBS was used to wash the primary antibodies and the cryosections were incubated with secondary antibodies in PBS with the PBS containing 2% NDS and 0.1% Triton X-100 for 1 hour. The following primary antibodies were used for staining: anti-FOXG1 (rabbit, Takara, M227, 1:400 dilution), anti-TCF7L2 (rabbit, Cell Signaling Technology, 2569S, 1:200 dilution), anti-PHOX2A (rabbit, Abcam, ab155084, 1:200 dilution), anti-BRN3A (mouse, Sigma, MAB1585, 1:200 dilution), anti-mCherry (goat, Biorbyt, orb153320, 1:1000 dilution), anti-NK1R (rabbit, Sigma, S8305, 1:200 dilution). Alexa Fluor dyes (Life Technologies) were used at 1:200 to 1:1,000 dilution and nuclei were visualized with the Hoechst 33258 dye (Life Technologies, H3549, 1:10,000 dilution). Cryosections were mounted for microscopy on glass slides using Aqua-Poly/Mount (Polysciences, 18606) or Vectashield (Vector labs, H-1000), and imaged on a confocal microscope. Images were processed and analyzed using Fiji (ver. 2.14.0) and IMARIS (Oxford Instruments).

### Design and 3D printing of custom wells for loop assembly

The custom wells for the loop assembloids were designed using Autodesk 3ds Max (Autodesk). The dimension and spacing between the individual parts were tailored to accommodate the size of the organoids. The wells were 3D-printed using the Original Prusa i3 MK3S + 3D printer (Prusa Research) with a biocompatible polyester polylactic acid (PLA) thermoplastic filament (Prusa Research). The printed structures were filled with 100% infill and randomized seams.

### Single cell RNA-seq library preparation and data analysis

Dissociation of regionalized neural organoids was performed as described previously ^49,50^. Briefly, 4–5 randomly selected organoids were pooled to obtain single cell suspension from regionalized organoids, and then incubated in 30 U/mL papain enzyme solution (Worthington Biochemical, LS003126) and 0.4% DNase (12,500 U/mL, Worthington Biochemical, LS2007) at 37 °C for 45 minutes. After enzymatic dissociation, spheres were washed with a solution including protease inhibitor and gently triturated to achieve a single-cell suspension. Cells were resuspended in 0.04% BSA/PBS (MilliporeSigma, B6917-25MG) and filtered through a 70 μm Flowmi Cell Strainer (Bel-Art, H13680-0070), and the number of cells were counted. To target 7,000 cells after recovery, approximately 11,600 cells were loaded per lane on a Chromium Single Cell 3′chip (Chromium Next GEM Chip G Single Cell Kit, 10x Genomics, PN-1000127). cDNA libraries were generated with the Chromium Next GEM Single Cell 3′ GEM, Library & Gel Bead Kit v3.1 (10x Genomics, PN-1000128), according to the manufacturer’s instructions. Each library was sequenced using the Illumina NovaSeq S4 2 × 150 bp by Admera Health. Quality control, UMI counting of Ensembl genes and aggregation of samples were performed by the ‘count’ and ‘aggr’ functions in Cell Ranger software (version, Cellranger-7.0.1). Further downstream analyses were performed using the R package Seurat (version 5.0.1) ^51^.

To ensure only high-quality cells were included for downstream analyses, an iterative filtering process was implemented for each sample. First, cells with less than 2,000 unique genes and with mitochondrial counts greater than 10% of the total counts were identified and removed. Subsequently, raw gene count matrices were normalized by regularized negative binomial regression using the sctransform function (vst.flavor=“v2”), which also identified the top 3,000 highly variable genes using default parameters. Dimensionality reduction using principal component analysis (PCA) on the top variable genes was performed and clusters of cells were identified in PCA space by shared nearest-neighbor graph construction and modularity detection implemented by the FindNeighbors and FindClusters functions using a dataset dimension of 50 (dims=50 chosen based on visual inspection of elbow plot) with default parameters. We subsequently performed iterative rounds of clustering (resolution = 2) to identify and remove clusters of putative low-quality cells based on outlier low gene counts (median below the 10th percentile), outlier high fraction mitochondrial genes (median above the 95th percentile), and/or high proportions of putative doublets identified by the DoubletFinder package ^52^ (median DoubletFinder score above the 95th percentile). We iteratively removed low-quality clusters until no further clusters met the above thresholds. Following low quality cell removal, an additional round of filtering of putative stressed cells was performed inspired by the Gruffi R package workflow ^53^. Specifically, putative clusters that contained high module score expression (z-score > 1) of endoplasmic reticulum stress signature genes (defined by GO:0034976 gene set and calculated using AddModuleScore function) were identified and removed. All individual 10x samples (n = 12) were integrated using the IntegrateData function with the above parameters using the reciprocal PCA method. The integrated dataset was clustered (FindClusters function; resolution = 1) and embedded for visualization purposes with Uniform Manifold Approximation and Projection (UMAP). We identified and categorized major cell classes through a combination of marker gene expression and annotation via reference mapping using Seurat’s TransferData workflow to classify organoid cells based on a well-annotated human fetal scRNA-seq dataset ^32^. GABAergic neurons were identified (*GAD1* and *GAD2* containing clusters) and re-processed and subclustered (FindClusters function; resolution = 1). Marker genes for each clustered were identified using the FindMarkers function with default parameters.

### Viral labeling and live cell imaging

Viral infection of 3D neural organoids was performed as previously described ^29^. Briefly, regionalized neural organoids (1-3 organoids per one Eppendorf tube) were transferred into a 1.5 mL Eppendorf tube containing 200 μL of the neural medium, and incubated with the virus overnight at 37°C, 5% CO_2_. The next day, fresh 800 μL of culture media was added. The following day regionalized neural organoids were transferred into fresh culture media in ultra-low attachment plates (Corning, 3471, 3261). The viruses used in this study were: AAV-DJ-hSYN1::EYFP (Stanford University Neuroscience Gene Vector and Virus Core, GVVC-AAV-16), AAV-DJ-hSYN1::mCherry (Stanford University Neuroscience Gene Vector and Virus Core, GVVC-AAV-17), AAV-DJ-EF1α::CVS-G (Produced by Stanford University Neuroscience Gene Vector and Virus Core using Addgene, plasmid #67528)^54^, Rabies-ΔG-Cre-GFP (Salk), AAV-DJ-EF1α-DIO-tdTomato (Stanford University Neuroscience Gene Vector and Virus Core, GVVC-AAV-169), AAV-EF1a-DIO-oG (Produced by Stanford University Neuroscience Gene Vector and Virus Core using Addgene, plasmid # 74290)^55^, Rabies-ΔG-Flpo-DsRedX (Salk, using Addgene, plasdmid #32650)^56^, AAV-CAG-fDIO-oG (Produced by Stanford University Neuroscience Gene Vector and Virus Core using Addgene, plasmid # 74291)^55^, Rabies-ΔG-TagBFP (Salk), AAV-DJ-hSYN1::mTurquoise2 (Addgene, #99125, AAV generated by Stanford University Neuroscience Gene Vector and Virus Core)^57^, and AAV-hSYN1::jGCaMP8s (Produced by Vectorbuilder)^35^.

### Generation of human loop assembloids

To generate human loop assembloids, hCO, hStrO, hMeO, and hDiO were separately generated from the same hiPS cells, and later assembled by placing them in close proximity with each other in a custom-designed 3D printed well for 4-7 days in a CO_2_ incubator. Formed assembloids were placed in 6-well ultra-low attachment plates in the neural medium described above using a cut P1000 pipette tip. After the assembly, the medium was changed every 4-7 days.

### Viral transsynaptic tracing in 2-or 4-part assembloids

For rabies virus retrograde tracing in 2-part assembloids, organoids representing the ‘presynaptic’ part were labeled with AAV-DIO-tdTomato, and organoids representing the ‘postsynaptic’ part were separately labeled with Rabies-ΔG-Cre-GFP and AAV-DJ-EF1α::CVS-G-WPRE-pGHpA. For 3-step rabies retrograde tracing in 4-part assembloids, hDiOs were infected with AAV-DJ-EF1α::CVS-G-WPRE-pGHpA and ΔG-Rabies-Cre-GFP, hMeOs were infected with AAV-DJ-EF1α::DIO-oG and ΔG-Rabies-Flpo-DsRedX, hStrOs were infected with AAV-DJ-CAG::fDIO-oG and ΔG-Rabies-TagBFP. Two days after the viral infection and washing out viruses, organoids were assembled. After 3 weeks of fusion, assembloids were imaged with a Leica TCS SP8 or Leica Stellaris 5 confocal microscope and analyzed in Fiji (ImageJ, version 2.1.0, NIH).

### Volumetric and mesoscale calcium imaging of assembloids

Assembloids expressing jGCaMP8s were imaged using a custom volumetric and mesoscale functional imaging system (VoluScan™; VSFM_E(460-490)_CAM(500-550), Doric Lenses) with VoluScan™ Driver (VSD, Doric Lenses) and VoluScan™ multi-plane disk (4-planes, 100 μm Z-stack resolution, VSMD-4P 0-100-200-300, Doric Lenses), which allows fast and multi-Z-plane functional imaging (5 Hz for 4 planes with z-stack resolution at 100 μm) with a wide field of view (FOV; the theoretical FOV is 5.063 × 3.164 mm for 1920 × 1200 pixel) at the single-cell level (the theoretical pixel size; 2.637 μm x 2.637 μm). More specifically, a low-magnification objective lens (5x, N.A. 0.16, Olympus) is combined with a 100 mm focal length tube lens to get a magnification of 2.22x and capture a large FOV with minimal distortion. For imaging jGCaMP8s, the light illumination was provided using a LED light source at 475 nm (CLED_HP_475, Doric Lenses) controlled with Doric Neuroscience Studio Software (Version 6, Doric Lenses), and passed through an illumination collimator followed by a band-pass excitation filter (460-490 nm), while the detection path includes a band-pass emission filter (500-550 nm) and directly projects onto the CMOS camera (DMK 33UX174, The Imaging Source). Assembloids were transferred into a 24-well glass bottom plate (P24-0-N, Cellvis) and the plate was placed inside the Tokai Hit stage top incubator (TK-STXG-WSKMX-SET, Tokai Hit) on top of the widefield imaging system. The samples were incubated at 37°C with 5% CO_2_ for 15 min before imaging acquisition, and imaging was performed in an environmentally controlled condition (37°C with 5% CO_2_) with a 5 Hz acquisition speed using Doric Neuroscience Studio (Version 6, Doric Lenses). Raw image files including 4 different z-planes were once exported to TIFF files and interleaved to a single file using a custom code. Suite2p GUI^36^ on Python was used to automatically and manually select a region of interest (ROI) in the 4-plane images. The correlation between calcium traces was quantified using scaled correlation analysis (SCA)^58^. Traces were first averaged over 1-second intervals and partitioned into 20-second-long segments. The Pearson correlation coefficient (r) was computed between corresponding segments and averaged across all segments. This analysis was repeated for multiple time lags within the range of [-20, 20] seconds for each pair of calcium traces, with 1 second shift step. The peak correlation value across time lags was taken as a measure of synchrony between the calcium signals. Shuffled data for comparison were generated by randomly permuting the averaged 1-second segments within each calcium trace.

### Extracellular recording and analyses

Assembloids were embedded into 3% of low-melting gel agarose (IBI Scientific, IB70056) and transferred into artificial cerebrospinal fluid (aCSF) containing 124 mM NaCl, 3 mM KCl, 1.25 mM NaH_2_PO_4_, 1.2 mM MgSO_4_, 1.5 mM CaCl_2_, 26 mM NaHCO_3_ and 10 mM *D*-(+)-glucose with addition of GlutaMAX (Gibco, 35050061). Embedded organoids or assembloids were placed on Brain Slice Chamber-2 (Scientific Systems Design Inc, S-BSC2) and perfused with aCSF (bubbled with 95% O_2_ and 5% CO_2_) at 37°C; temperature was controlled and retained at 37°C by connecting to Proportional Temperature Controller PTC03 (AutoMate Scientific, S-PTC03). Acute 32channel P-1 probe with 2 shanks (CAMBRIDGE NeuroTech, ASSY-37 P-1) were connected to Acute probe adaptor; 32 channels Samtec to Omnetics (CAMBRIDGE NeuroTech, ADPT A32-Om32). Signals were acquired using the RHD recording controller (Intan Technologies), at 30000 Hz.

Raw recording data (in .rhd format) were opened using the Intan Technologies code and processed in Python. Single unit activity was extracted from the raw extracellular recordings using the Kilosort 4^59^ (version 4.03) with the Spikeinterface package^60^ (version 0.100.5). Samples showing large noise or aberrant signals distorting further analysis processes were excluded from further analysis. Following single unit extraction, units were assigned to specific organoids based on their spatial locations on the recording probes. For spontaneous firing rate analysis, the total number of spikes was divided by the recording duration. For the scaled correlation analysis (SCA), spike trains were binned into 1-second bins, and correlations were computed on a time scale of 20 seconds. Specifically, the SCA splits the binned spike trains into consecutive 20-second segments, calculating the Pearson correlation coefficient (r) between co-occurring segments, with 0.5 second shift step. The overall correlation between a pair of spike trains is given by the average of r values across all segments. This analysis is repeated across different time lags within the range [-20, 20] seconds, and the maximal correlation value is used as the measure of spike train synchrony.

### Genome editing of hiPS cells to generate ASH1L+/-hiPS cell lines

Synthetic sgRNA was synthesized and purchased from Synthego with 2′-O-methyl 3′-phosphorothioate modified nucleotides at the three terminal positions at both the 5′ and 3′ ends. The genomic target sequences of the sgRNAs were 5’-gcaacgccatctactcttct-3’. hiPS cells were washed with DPBS and incubated with 1 mL Accutase at 37 °C for 10 min, after which 9 mL of Essential 8 medium was added to the well for resuspension. A total of 3 × 10^6^ cells were centrifuged and nucleofected using the P3 Primary Cell 4D-Nucleofector X Kit L (Lonza, V4XP-3024), a 4D-nucleofector core unit, and an X unit (Lonza) with the DC100 program. After nucleofection, cells were resuspended with the warm medium in the nucleofection cuvette and seeded into a well of a 6-well plate containing a pre-warmed Essential 8 medium supplemented with the ROCK inhibitor Y-27632. At 2–3 days after nucleofection, cells were dissociated with accutase and GFP^+^ cells were sorted into 96-well plates using a BD Aria II (Stanford Shared FACS Facility). Sorted single cells were cultured in StemFlex medium (ThermoFisher Scientific, A334940) and 10 U/mL Penicillin-Streptomycin (Gibco, 15140122) with every other day half media changes for single colony formation in individual wells. After 2-3 weeks, colonies were passaged into 96-well plates, and isolated clones were expanded, cryopreserved, and collected for genotyping analysis.

The genotype of the genome-edited clones was determined by PCR with the primer set FW, 5’-GCCTCTCGTATAGCAGCAGA-3’; RV, 5’-ACAGAGCCAACACCTGCTTT-3’, and Sanger sequencing was performed (Genewiz) with the same primers. ICE analysis (https://ice.synthego.com/) was performed on obtained sequencing chromatograms to determine the genotype. The genotyping consists of Sanger sequencing and next-generation sequencing (NGS) on the amplicons encompassing the gRNA-targeting region. For Sanger sequencing, Phire Tissue Direct PCR Master Mix (ThermoFisher Scientific, F170L) was used according to the manufacturer’s recommendations, while the GoTaq Flexi DNA Polymerase (Promega, M7805) was used for NGS analysis. DNA was isolated using the DNeasy Blood & Tissue Kit (Qiagen, 69506). An unmodified hiPS cell lines line that underwent the same CRISPR editing process but was not genetically modified was selected as a control/isogenic hiPS cell line. After genome editing, SNP arrays were performed for isogenic and *ASH1L*^+/-^ lines.

### Statistics

Data are presented as Mean ± s.e.m. Raw data were tested for normality of distribution, and statistical analyses were performed using unpaired t-test (two-tailed), Mann-Whitney test, Wilcoxon matched-pairs signed rank test or Kruskal-Wallis test with Dunn’s multiple comparisons test depending on the dataset and experimental designs. Sample sizes were estimated empirically. GraphPad Prism Version 9.3.1 was used for statistical analyses.

### Data and materials availability

Gene expression data were deposited in the Gene Expression Omnibus under accession number GSE279264. The codes used for calcium imaging and electrophysiology analyses in this study have been deposited at Zenodo (DOI: 10.5281/zenodo.13921410).

**Supplementary Data Figure 1.**
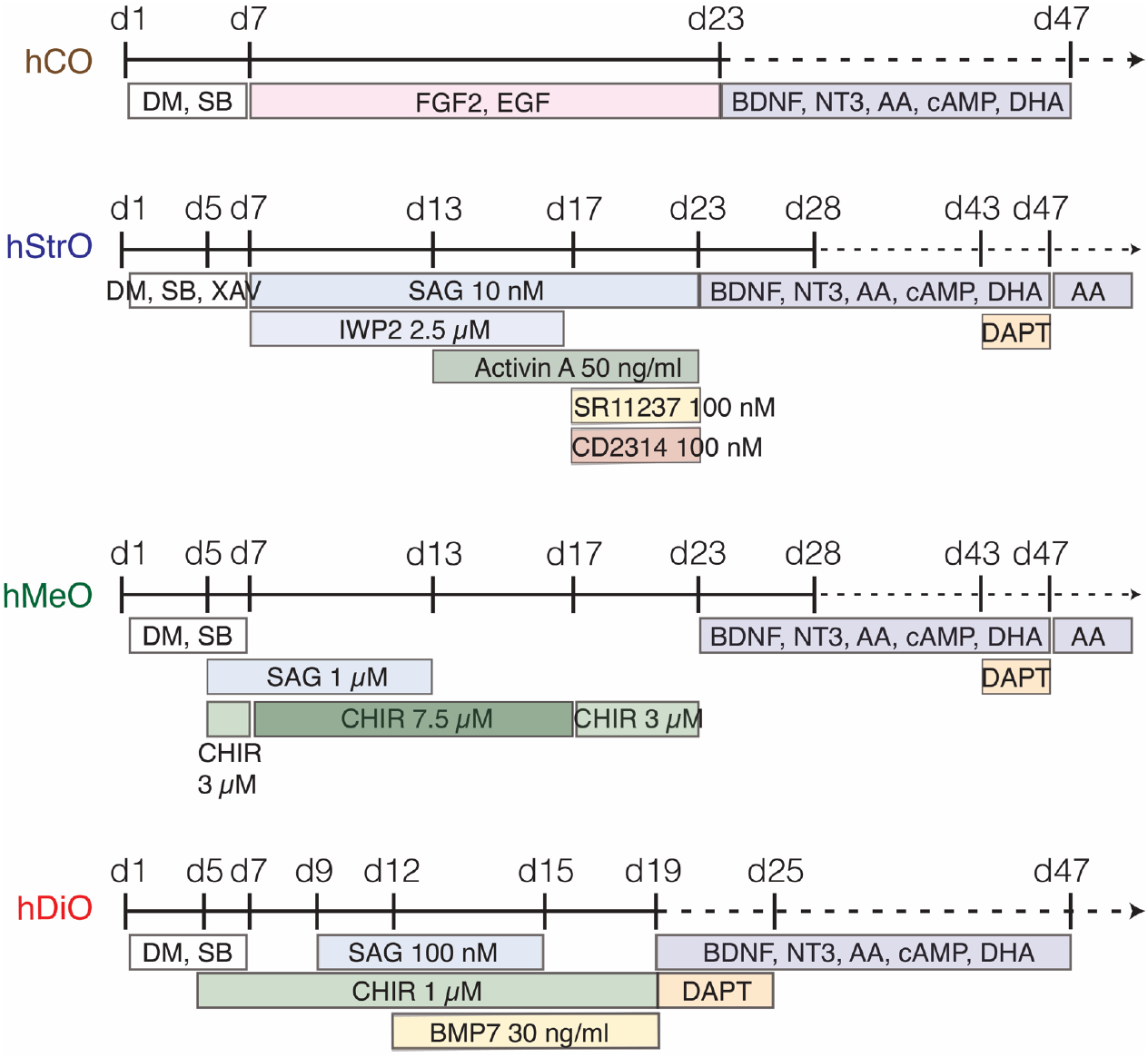
Protocol for generating regionalized neural organoids. Protocols to generate hCO, hStrO, hMeO, and hDiO from hiPS cells.

**Supplementary Data Figure 2.**
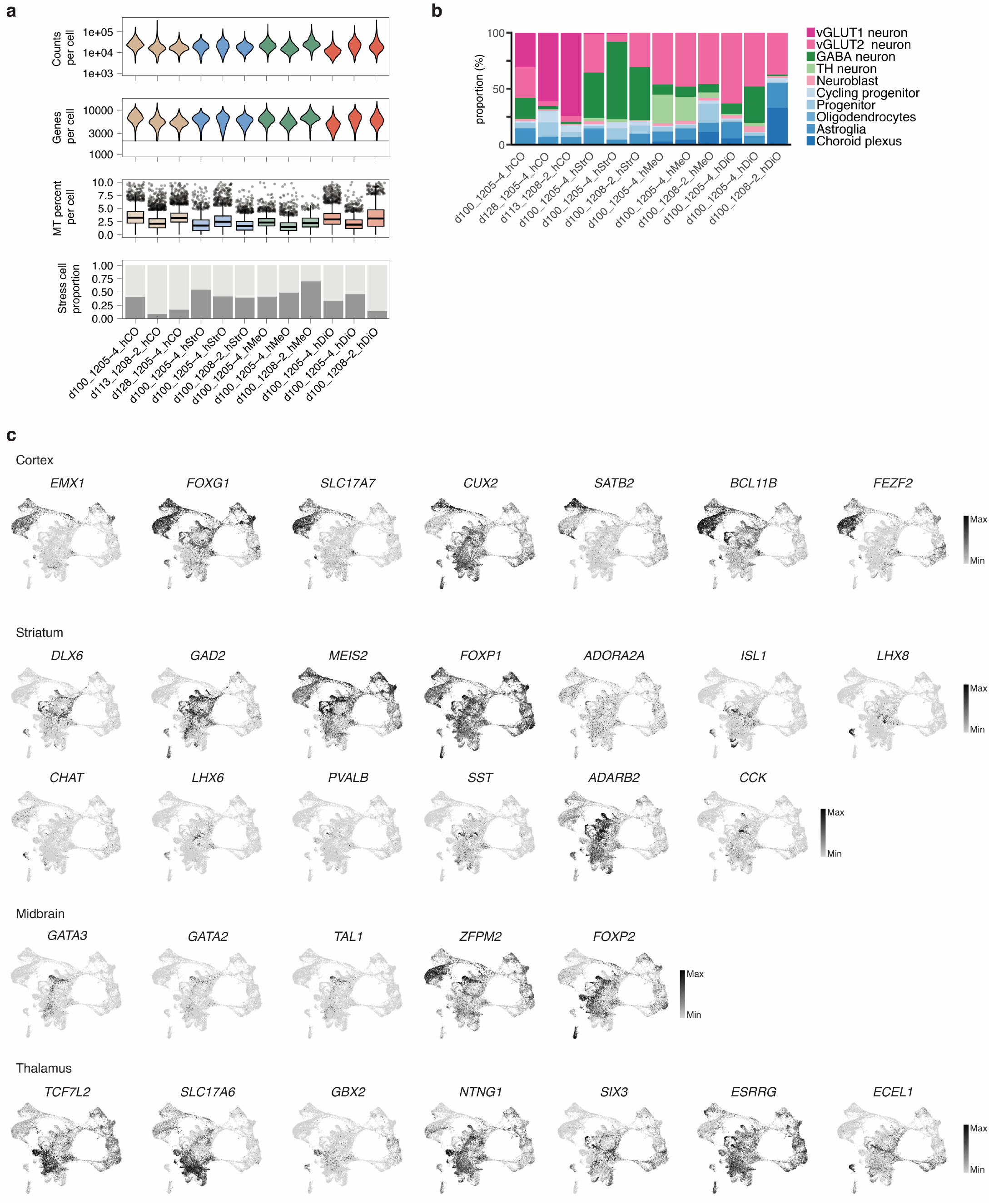
Single cell transcriptomics of regionalized organoids. **a**. Plots for scRNA-seq quality showing the distribution of the number of counts per cell, number of genes per cell, mitochondrial (MT) gene percentage per cell, and stress cell proportion in each sample. Mitochondrial gene fraction plotted as boxplots (horizontal line denotes median; lower and upper hinges correspond to the first and third quartiles; whiskers extend 1.5 times the interquartile range with outliers shown outside this range). Lines denote cell quality thresholds. **b**. Distribution of cell clusters across each sample. **c**. UMAP visualization of expression for cell type marker genes.

**Supplementary Data Figure 3.**
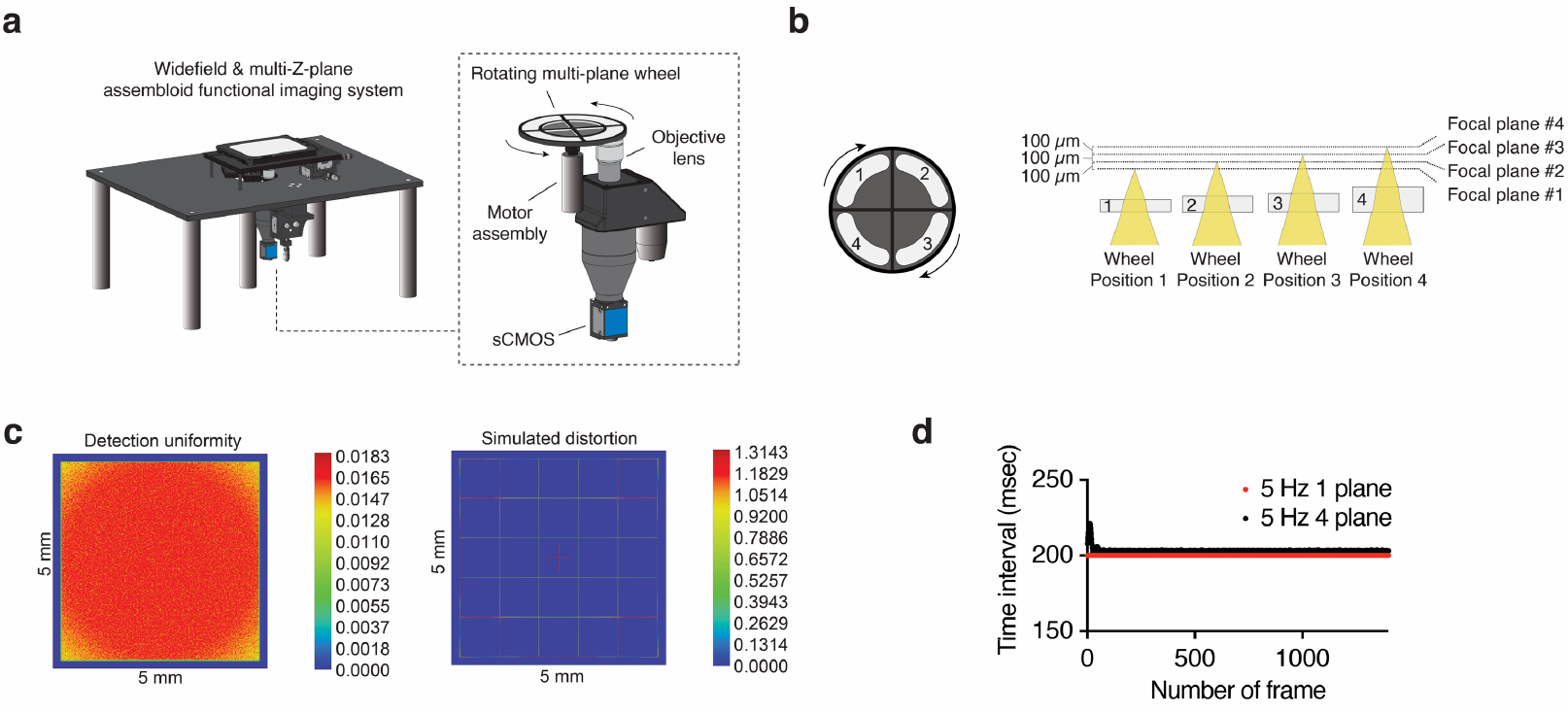
Multi-Z-plane imaging set up. **a**. Schematic illustrating a widefield and multi-Z-plane assembloid functional imaging system with temperature and CO2 level regulation chamber. Inset highlights that a multi-Z-plane wheel is rotating on top of the objective lens during imaging acquisition. **b**. Schematic illustrating a rotating multi-Z-plane wheel with different thicknesses of windows (left) and four focal planes imaging with 100 μm distance between planes, determined by different wheel positions. **c**. Consistency of detection uniformity with minimal simulated distortion values across the entire wide field of view. **d**. Consistency of time interval during imaging acquisition over one thousand frames with multiple Z-plane imaging.

**Supplementary Data Figure 4.**
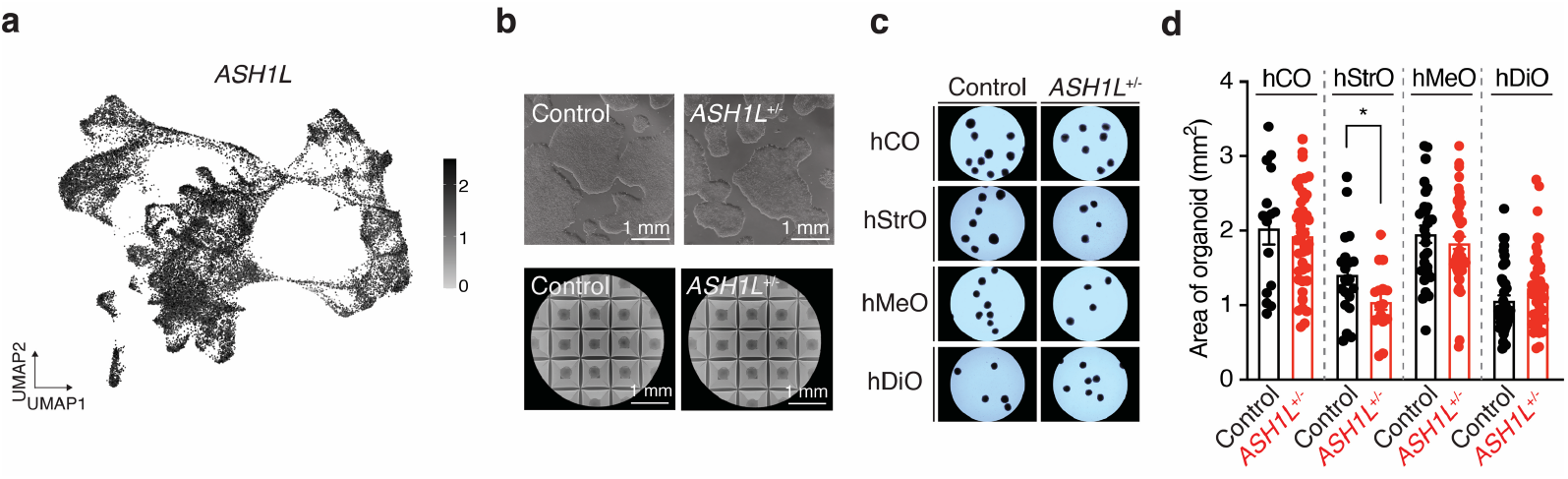
Characterization of *ASH1L*^**+/-**^ hiPS cells and organoids. **a**. UMAP plot showing *ASH1L* expression patterns in the neural organoids in hLA. **b**. Representative images showing the colony of control and *ASH1L*^+/-^ hiPS cells (top) and cell formation of 3D regionalized organoids (bottom). **c**. Representative images of 3D neural organoids from control and *ASH1L*^+/-^ hiPS cell lines. **d**. Quantification of the size of four neural organoids from control and *ASH1L*^+/-^ hiPS cell lines (hCO; n = 15 organoids for control, n = 42 organoids for *ASH1L*^+/-^ / hStrO; n = 20 organoids for control, n = 16 organoids for *ASH1L*^+/-^ / hMeO; n = 29 organoids for control, n = 47 organoids for *ASH1L*^+/-^ / hDiO; n = 43 organoids for control, n = 40 organoids for *ASH1L*^+/-^ from 3 differentiations, Unpaired t-test test, \**P* = 0.0471).

**Supplementary Data Figure 5.**
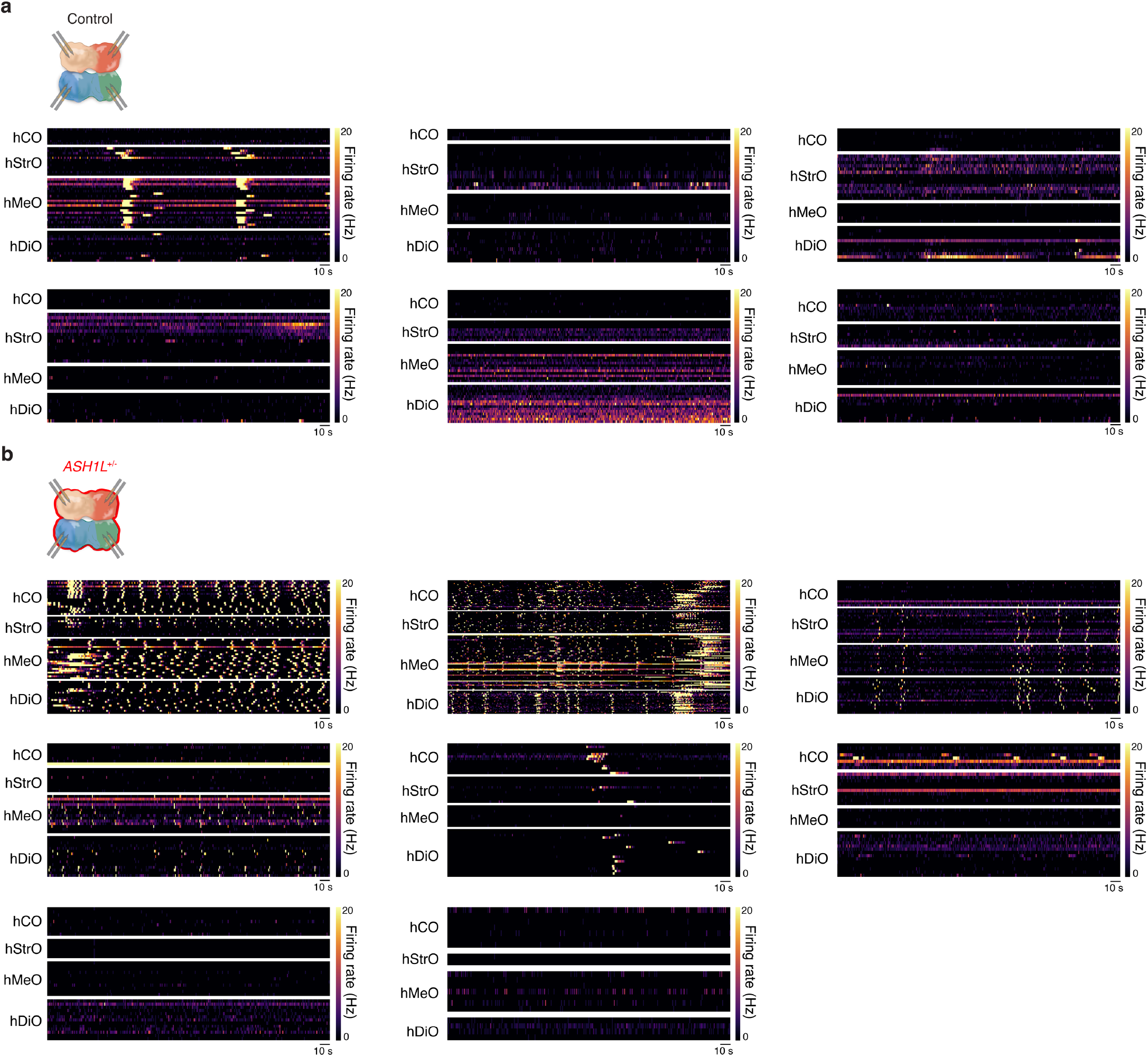
Activity heatmaps of control and *ASH1L*^+/-^human loop assembloids. **a**. Schematics illustrating four region extracellular recordings of control hLA and activity heatmaps of all 6 samples used for **Figure 5**. **b**. Schematics illustrating four region extracellular recordings of *ASH1L*^+/-^ hLA and activity heatmaps of all 8 samples used for **Figure 5**.

**Supplementary Table 1. scRNA-seq cluster annotation and marker genes.**

Marker genes for each annotated cell cluster as determined using the Seurat FindMarkers function with default parameters (two-sided Wilcoxon Rank Sum test with Bonferroni correction).

**Supplementary Table 2. GABAergic scRNA-seq subcluster annotation and marker genes**

Marker genes for each annotated cell cluster as determined using the Seurat FindMarkers function with default parameters (two-sided Wilcoxon Rank Sum test with Bonferroni correction).

